# Indirect genomic effects shape cancer risk across species

**DOI:** 10.64898/2026.06.29.735167

**Authors:** George Butler, Srividya Ramakrishnan, Tyler Collins, Joanna Baker, Sarah R. Amend, The Vertebrate Genomes Project Consortium Phase I, Michael C. Schatz, Chris Venditti, Kenneth J. Pienta

**Author notes:** Corresponding authors: George Butler, Michael C. Schatz, Chris Venditti.

## Abstract

Tumour prevalence varies dramatically throughout the animal kingdom despite broadly conserved cellular and developmental processes, raising the question of how evolution has shaped susceptibility ^1,2^. Here, we link macroevolutionary variation in tumour prevalence to gene-level selection by integrating comparative genomics data from 109 species of birds and mammals using a Bayesian phylogenetic framework to estimate pangenome-wide rates of genetic evolution across >150 million years of evolutionary change. We identify 3,206 genes in which natural selection is associated with shifts in tumour prevalence, with more than 80% of which are linked to reduced prevalence, suggesting pervasive selection for cancer suppression. Using causal phylogenetic inference, we show that genes associated with reduced tumour prevalence act predominantly through indirect effects on body size, revealing growth as a key mediator of cancer risk across species. In contrast, genes associated with increased tumour prevalence exert direct effects independent of body size. Finally, at the species-level, we demonstrate that exceptionally low rates of benign tumours do not necessarily coincide with reduced malignancy, revealing that benign and malignant tumour processes are evolutionarily decoupled. Together, these results reveal how natural selection has fine-tuned the link between genotype, phenotype, and cancer risk across species.

## Main

The evolutionary relationship between body size and cancer illustrates a central question in biology – how genetic changes translate into phenotypic outcomes ^3,4^. The genomic mechanisms that lead to diversity in species size, including accurate DNA replication, tightly controlled cellular proliferation, and long-term genomic stability, are the same processes that break down and give rise to tumours ^5–7^. Across terrestrial vertebrates, larger species tend to develop more tumours than smaller species ^2,8,9^, illustrating how growth and oncogenesis can emerge from the same molecular foundation.

Previous comparative genomics studies have characterised how tumour prevalence varies with gene paralogs ^10^, copy number changes ^11,12^, accelerated genomic regions ^13^, gene conservation ^14^ and immune surveillance ^15^. However, recent evidence has shown that while tumour prevalence increases with body size, species that evolved larger bodies more quickly have a lower prevalence of malignant tumours than expected for their size ^8,9^. This suggests that natural selection has repeatedly favoured mechanisms that allow these species to grow larger while mitigating the risk of lethal cancer ^16^. However, the molecular basis of such trade-offs is unknown. For example, does selection act on individual genes to reduce tumour prevalence directly, indirectly through effects on body size (or body size-associated phenotypes), or through a combination of both processes (Figure 1A)? Identifying genes that shape tumour prevalence directly, independent of body size, will increase our understanding of cancer progression and may provide insight for the design of therapeutic strategies to target malignant cells without impacting normal proliferative tissue.

**Figure 1.**
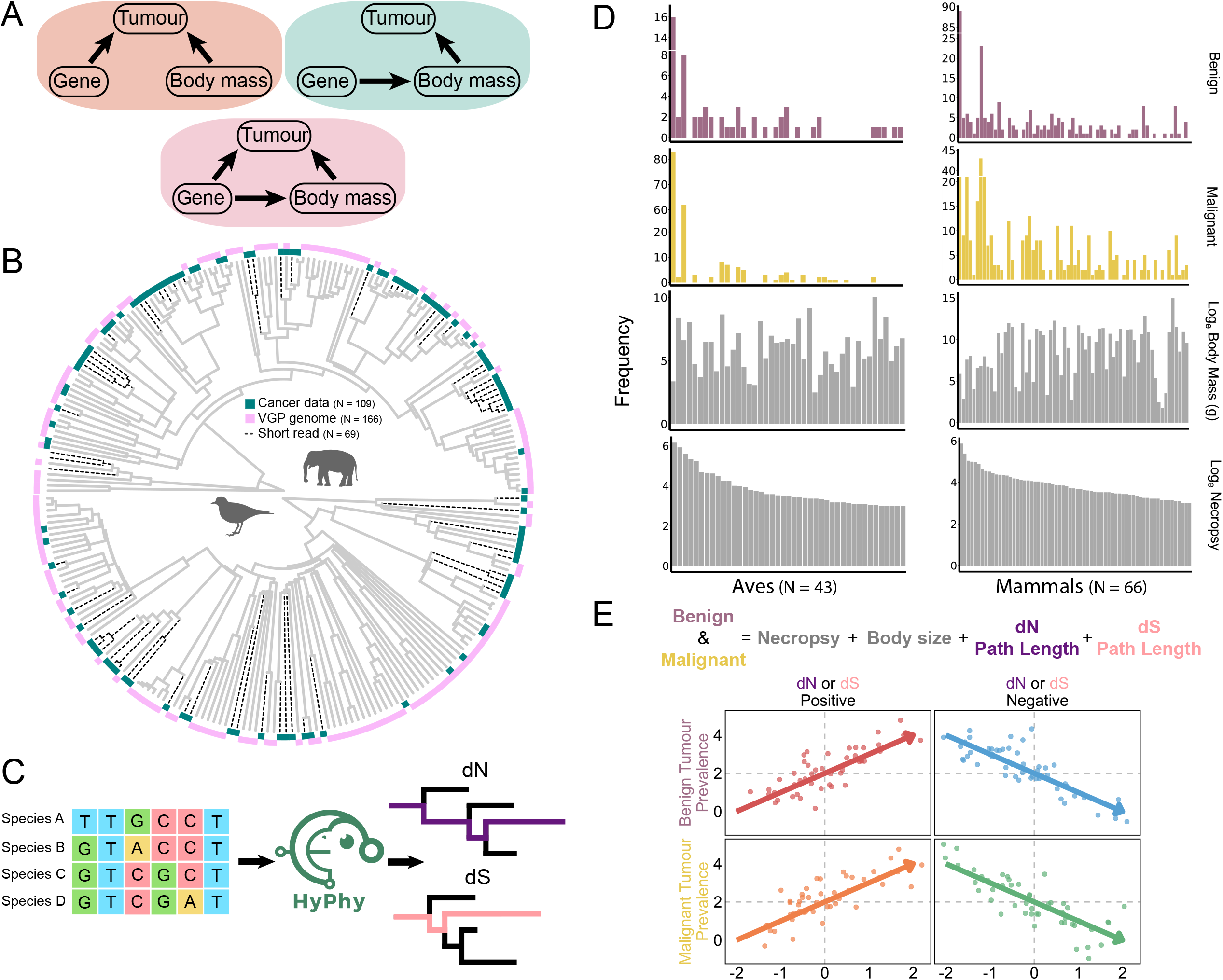
Measuring molecular adaptation. **A**) An illustration of a direct relationship between tumour prevalence and gene evolution (orange), and an indirect relationship via body size (green), or a combination of the two (pink). **B**) A phylogenetic tree of the 256 bird and mammal genomes used to estimate rates of molecular evolution. Genomes from the VGP phase 1 release are shown at the tips of the tree (pink) as are species with paired cancer data (green). Species with genomes from short read sequencing are shown by branches with dotted lines. **C**) A schematic of the pipeline used to estimate species-specific rates of non-synonymous (dN) and synonymous (dS) evolution for each gene. Multiple sequence alignments (MSAs) are built for each gene, and an adaptive branch-site random effect likelihood (aBSREL) model is used to estimate separate non-synonymous and synonymous rate-scaled phylogenetic trees. The species-specific amount of non-synonymous and synonymous evolution is measured as the distance from root to tip. **D**) Histograms showing the distribution of benign and malignant tumour prevalence, body size, number of genes analysed, and number of necropsies. E) The fixed effects of the multivariate phylogenetic generalised linear mixed model (MPGLMM) used to test for an association between tumour prevalence and gene evolution. The potential directionality of the four different significant benign or malignant associations with dN or dS are shown: positive association between benign prevalence and dN or dS (red), negative association between benign prevalence and dN or dS (blue), positive association between malignant prevalence and dN or dS (orange), and negative association between malignant prevalence and dN or dS (green).

Here, we use high-quality reference genomes from the Vertebrate Genomes Project (VGP) ^17^, complemented with genome sequences from other projects, with tumour prevalence and body size data across 109 species of birds and mammals within a Bayesian phylogenetic framework to unravel how natural selection has balanced the evolution of body size and tumour prevalence. Estimating species-specific protein altering and non-altering changes (non-synonymous and synonymous substitutions) across more than 14,000 individual genes, we test how evolutionary rates predict benign and malignant tumour prevalence while accounting for body size, sampling, and shared ancestry. We then use evolutionary causal modelling to determine the relative proportion of genes that influence tumour prevalence directly, indirectly through body size, or through both processes simultaneously. This framework enables a systematic dissection of the molecular mechanisms underlying the co-evolution of body size and cancer risk across species. Together, these analyses show that genes associated with tumour prevalence operate through both body size-dependent and body size-independent pathways, while also revealing that benign and malignant tumour processes have distinct evolutionary architectures.

### Measuring the strength of molecular adaptations

To investigate how individual genes have shaped the co-evolution of cancer prevalence and body size among birds and mammals, we used a Bayesian multivariate phylogenetic generalized linear mixed model (MPGLMM) framework (Methods) ^18–20^. Within this framework, we examined the genomes from 256 bird and mammal species, including 166 from VGP ^17^, along with an additional 90 from other large-scale genome sequencing projects (Supplementary Table S1) matched with cancer prevalence data from 109 species^2^. While most genomes were high-quality long-read assemblies, a limited number of short-read assemblies were retained to maximize cancer prevalence coverage (N = 69, Supplementary Table S1). Specifically, assemblies submitted before 2015, contig-scale assemblies, and scaffold-scale genomes with available chromosome-scale versions were excluded, with only the most recent assembly retained per species. To systematically identify orthologs across these diverse species, we implemented an optimized alignment-based strategy using Miniprot ^21^ to annotate 20,421 reviewed human UniProt ^22^ genes (Methods, Supplementary Table S1-S2). Genes were retained if represented in at least 50 species and accompanied by an amphibian outgroup, yielding 15,124 genes for downstream analysis. On average, 10,672 genes were successfully recovered per species (range: 1,114 – 19,435; median 8105). For each retained gene, multiple sequence alignments (MSAs) were constructed using the OrthoMAM v12 pipeline ^23^, incorporating species with paired tumour data alongside an additional 163 high-quality genomes from the VGP to improve molecular rate estimation (Figure 1B). Each alignment contained a mean of 188 species (range: 50-280; median: 144), and both identity and positive substitution scores consistently exceeded the 0.75 retention threshold across both bird and mammal genomes, confirming high alignment quality (Supplementary Figure 1).

Variation in the rate of molecular evolution was estimated for each gene by fitting an adaptive branch-site random effect likelihood (aBSREL) model to produce separate non-synonymous and synonymous rate-scaled trees ^24^. In turn, the amounts of species-specific non-synonymous evolution (dN) and synonymous evolution (dS), were measured as the root-to-tip distance for each species in each of the rate-scaled trees (Methods, Figure 1C) ^20^. We interpret dN as a measure of adaptive evolution through protein-coding evolutionary change and dS as nearly neutral evolution reflecting background molecular evolutionary processes ^25,26^. Benign (non-cancerous) and malignant (cancerous) tumour prevalence were modelled as a function of body size, dN, and dS, while controlling for the number of necropsies per species and the shared ancestry, henceforth referred to as the full model (Figure 1D & E, see methods). Throughout, we refer to dN-positive or dN-negative genes (and similarly dS) to represent the directionality of significant associations between the covariate and tumour prevalence (Figure 1E). The final dataset consisted of 109 species, composed of 43 birds and 66 mammals, spanning 14,189 individual genes.

### Genome-wide patterns of gene evolution and tumour prevalence

We discovered strong selective pressures acting on malignant tumour suppression. Overall, 3206 genes had a significant association between tumour prevalence and dN (Figure 2A, Supplementary Table S3) with significantly more dN-negative (N = 2666) compared to dN-positive (N = 540) 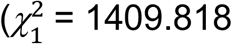, P < 0.001, N=3206). That is, where selection is acting to modify protein sequences, it tends to reduce the prevalence of cancer. In addition, a significantly higher proportion of dN-negative genes were associated with malignant compared to benign tumour prevalence (1421 compared to 701, 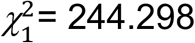, P < 0.001, N=2122). The effect size for genes is higher for malignancy than benign for both dN-negative (*t*_2705.946_ = 8.886, P < 0.001, *µ*_*Benign*_ = −0.488, *µ*_*Malignant*_ = −0.566) and dN-positive (*t*_560.926_ = 4.423, P < 0.001, *µ*_*Benign*_ = 0.421, *µ*_*Malignant*_ = 0.493) genes (Figure 2C, Extended Figure 1A & 1C). That is, the same level of non-synonymous evolution has a greater impact on malignancy compared to benign tumour prevalence.

**Figure 2.**
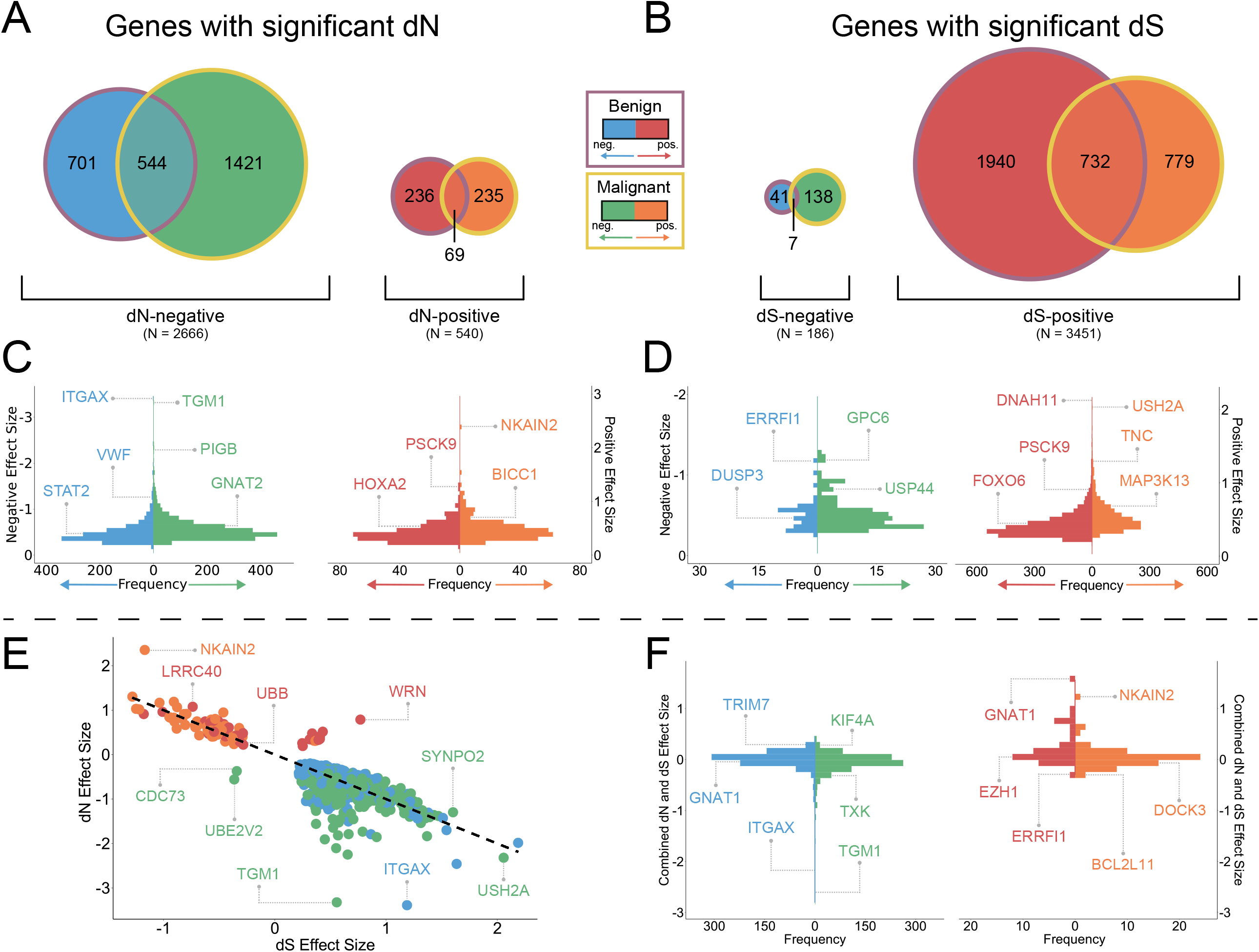
Genome-wide patterns of molecular evolution associated with tumour prevalence. **A**-**B)** The number of genes with a significant association between benign (blue or red) or malignant (green or orange) tumour prevalence and **A**) dN or **B**) dS. **A**) Significantly more genes have a negative (dN-negative, N = 2666) compared to positive (dN-positive, N = 540) association between tumour prevalence and dN 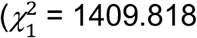, P < 0.001, N=3206), and significantly more dN-negative genes are associated with malignant compared to benign tumour prevalence (1421 compared to 701, 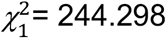, P < 0.001, N=2122). **B**) Significantly more genes have a positive (dS-positive, N=3451) compared to negative (dS-negative, N=186) association between tumour prevalence and dS 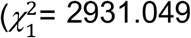, P < 0.001, N=3637), and significantly more dS-positive genes are associated with benign compared to malignant tumour prevalence (1940 compared to 779, 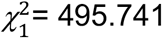, P < 0.001, N=2719). **C**-**D**) Distribution of standardized effect sizes for genes with a significant association between benign (blue or red) or malignant (green or orange) tumour prevalence and **C**) dN or **D**) dS. **C**) The effect size of dN-negative genes is significantly higher for malignancy compared to benign tumour prevalence (*t*_2705.946_ = 8.886, P < 0.001, *µ*_*Benign*_ = −0.488, *µ*_*Malignant*_ = −0.566) and for dN-positive genes (*t*_560.946_ = 4.423, P < 0.001, *µ*_*Benign*_ = 0.421, *µ*_*Malignant*_ = 0.493) **D**) The effect size for dS-positive genes is significantly higher for malignant compared to benign tumour prevalence (*t*_2704.500_ = 19.421, P < 0.001, *µ*_*Benign*_ = 0.424, *µ*_*Malignant*_ = 0.541). **E**) Relationship between dN and dS effect sizes for genes with significant effects for both covariates. Each point represents a gene, and the dashed line indicates the fitted linear regression. A significant negative association between the dN and dS effect size on tumour prevalence (β = −0.985, P < 0.001, N=1704, R^2^ = 0.697). **F**) The combined effect of dN and dS on tumour prevalence. The combined effect is significantly larger for dN-negative genes on malignant (green) compared to benign (blue) tumour prevalence (*t*_1475_ = 7.188, P < 0.001, *µ*_*Benign*_ = 0.003, *µ*_*Malignant*_ = −0.072) and significantly larger for dN-positive genes on benign (red) compared to malignant (orange) tumour prevalence (*t*_58.899_ = 3.079, P = 0.003, *µ*_*Benign*_ = 0.251, *µ*_*Malignant*_ = 0.056).

Of the 3637 genes identified as having a significant association between tumour prevalence and dS (Figure 2B, Supplementary Table S3), we found significantly more dS-positive genes (N=3451) compared to dS-negative (N=186) genes 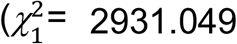, P < 0.001, N=3637). This suggests that higher rates of synonymous evolution derived from an increased occurrence of nearly neutral processes ^25,26^ is generally associated with higher tumour prevalence. Furthermore, significantly more dS-positive genes were associated with benign compared to malignant tumour prevalence (1940 compared to 779, 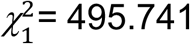, P < 0.001, N=2719) (Figure 2B), suggesting that benign, non-cancerous, tumour prevalence is more strongly shaped by neutral, background molecular evolutionary processes. However, akin to genes with significant dN, the magnitude of the effect for dS-positive genes was significantly higher for genes associated with malignant compared to benign tumour prevalence (*t*_2704.500_ = 19.421, P < 0.001, *µ*_*Benign*_ = 0.424, *µ*_*Malignant*_ = 0.541, Figure 2D, Extended Figure 1B & 1D).

To control for differences in species longevity, we also fitted a secondary model in which species longevity was included as an additional independent covariate (see Methods). We found qualitatively similar results between the two models suggesting that our findings were not being driven by differences in species longevity (Supplementary Figure 4).

To cross validate our results with human disease, we compared the overlap of dN and dS significant genes amongst the 14,189 genes in our dataset that are also found within three curated cancer gene datasets (see methods): COSMIC ^27^ (N=653), IntOGen ^28^ (N=544), and OncoKB ^29^ (N=1007). Across all three datasets, between 33%-35% of genes overlapped with genes with a significant dN or dS effect. Consistent with our genome-wide results, significantly more dN-associated genes, such as *MYC, RB1*, and *CCND1*, were associated with malignant prevalence (COSMIC 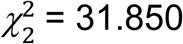, P < 0.001, N=120; IntOGen 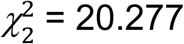, P < 0.001, N=101; OncoKB 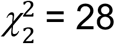, P < 0.001, N=206) (Extended Figure 1E) and significantly more dS-associated genes, such as *AKT1, MDM2*, and *FOXL2*, were associated with benign prevalence (COSMIC 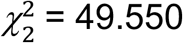, P < 0.001, N=160; IntOGen 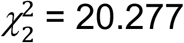, P < 0.001, N=132; OncoKB 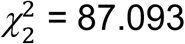, P < 0.001, N=247) Extended Figure 1F) within each dataset.

Across all significant genes, the relative contributions of dN and dS were further investigated by examining genes in which both covariates were significantly associated with either (or both) benign and malignant tumour prevalence (N = 1512 unique genes). Across these genes, there was a strong negative association between the effect sizes of dN and dS at the individual gene level (β = −0.985, P < 0.001, N=1704, R^2^ = 0.697, Figure 2E). That is, the stronger the positive dS effect promoting tumour evolution, the stronger the dN negative effect in reducing it. However, the slope of the dN and dS effect size was not significantly different from −1 (P = 0.334, N=1704), suggesting that an increase in tumour prevalence from dS is balanced by a corresponding decrease in tumour prevalence from dN. This antagonistic pattern may reflect a genome-wide evolutionary ‘tug-of-war’ in which processes associated with dS promote tumour susceptibility, whereas dN-associated processes counterbalance these effects, analogous to models of tumour evolution in which opposing forces shape progression ^30^. While this observation was generally consistent across genes, we also found a few exceptions. For example, *CDC73* and *UBE2V2*, (a tumour suppressor ^31^ and oncogene ^32^ respectively in human disease), have a negative effect on malignant tumour prevalence for both dN and dS.

Finally, the combined dN and dS effect shows further asymmetries. For dN-negative genes, the combined effect was significantly larger for malignant compared to benign tumour prevalence (*t*_1475_ = 7.188, P < 0.001, *µ*_*Benign*_ = 0.003, *µ*_*Malignant*_ = −0.072) (Figure 2F). In contrast, for dN-positive genes, the combined effect size was significantly larger for genes related to benign tumour prevalence (*t*_58.899_ = 3.079, P = 0.003, *µ*_*Benign*_ = 0.251, *µ*_*Malignant*_ = 0.056) (Figure 2F). Taken as a whole, these results show that either nearly neutral processes are weaker, or natural selection is stronger, for genes related to malignant tumour prevalence. This reinforces the notion that malignant tumours evolve under far stronger evolutionary constraints compared to benign tumours ^8^.

### Genomic influence on tumour prevalence and body size

To determine if the association between tumour prevalence, body size, dN, and dS arises through direct or indirect evolutionary relationships, we used a phylogenetic causal modelling framework ^33^. We first tested 11 different causal paths, representing all possible biologically plausible configurations of directional dependences between dN, dS, body size, and benign and malignant tumour prevalence (Extended Figure 2A). Eight of these paths met the criterion of conditional independence and thus were suitable for causal inference (Methods).

Across the eight eligible paths (Figure 3A), we found that the distribution of best-fitting paths differed strongly both between dN-positive and dN-negative genes, and between genes associated with benign and malignant tumour prevalence (Supplementary Table S4).

**Figure 3.**
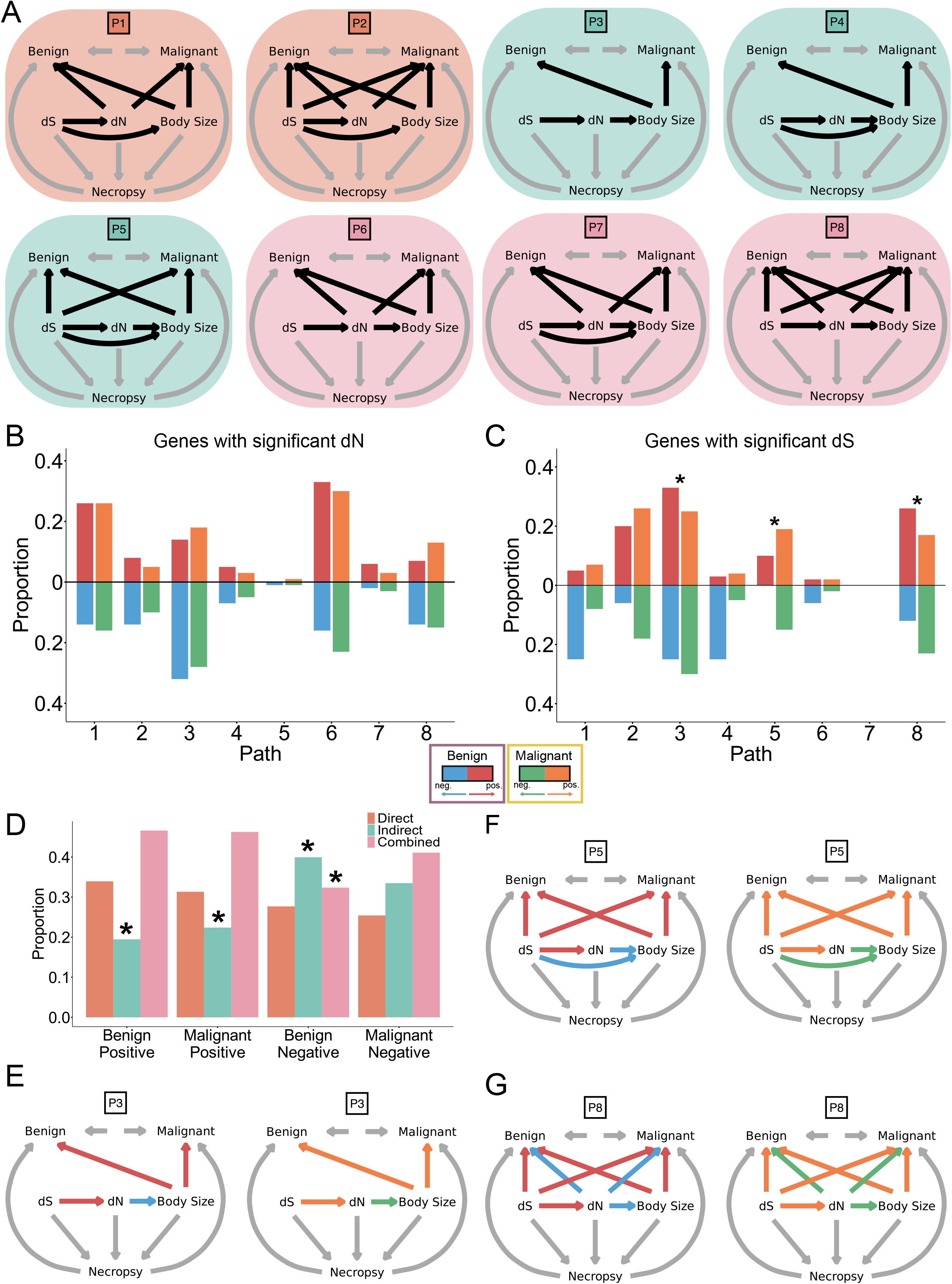
Causal relationships underlying tumour prevalence. **A**) A schematic of the eight different causal relationships (paths) tested between dS, dN, body size, and tumour prevalence. The direction of the arrow indicates the direction of the causal relationship. The background colour indicates whether the path is direct (orange; paths 1-2), indirect (green; paths 3-5), or combined (pink; paths 6-8). **B**-**C**) The proportion of genes with a significant **B**) dN or **C**) dS association stratified by the best fitting path. For dN-associated genes, indirect and combined paths were enriched among dN-negative genes, whereas direct and combined paths were more common among dN-positive genes. **C**) Paths 3 and 8 are significantly associated with benign tumour prevalence (P = 0.003 & P < 0.001 respectively) and path 5 is significantly associated with malignant tumour prevalence (P < 0.001). **D**) Summary of paths categories grouped into direct, indirect, and combined evolutionary relationships. Significantly less benign and malignant dN-positive genes are associated with an indirect path, and significantly more benign dN-negative genes are associated with indirect and combined paths. **E**-**G**) Representative examples of the most common causal architectures identified among genes with significant dS effects. In all cases, the colour of the arrows indicates whether there is an average positive (red or orange) or negative (blue or green) causal relationship as estimated from significant dS genes (**C**).

Among the dN-positive genes, path 6, representing a causal relationship from dS to dN, dN to body size and tumour prevalence, and body size to tumour prevalence, was the most common (N=133 out of 422) (Figure 3B). For many genes where increased dN corresponds to higher tumour prevalence, the effect of dN on tumour prevalence is both direct and mediated through body size. However, a direct effect of dN on tumour prevalence does not necessarily mean that the gene is not related to other life history traits, such as gestation or longevity. For example, in cancer, *SNAI1*, a gene in path 6, is associated with aggressive disease and worse outcomes in humans ^34^. In normal tissue, *SNAI1* plays a key role in wound healing ^35^, demonstrating that while the effect of a gene on tumour prevalence may be independent of body size, it may still have an important role in organismal function.

Among dN-negative genes, path 3, which represents a causal relationship from dS to dN, dN to body size, and body size to tumour prevalence, was the most common path (N = 368 out of 1276) (Figure 3B). This suggests that for most genes where increased dN corresponds to lower tumour prevalence, dN influences tumour prevalence indirectly via its influence on body size (or associated phenotypes).

Within genes with significant dS, we found, again, that path 3 was the most common irrespective of directionality (dS-positive, N = 693 out of 2281; dS-negative, N = 22 out of 77) (Figure 3C). Path 3 was also the most common among a randomly selected subset of genes with non-significant dN and dS effects (N = 224 out of 445) (Extended Figure 2B), suggesting that indirect, body-size mediated effects represent the prevailing causal architecture across the genome.

### Summarizing genomic influence on tumour prevalence

To provide a broader synthesis, we grouped each of the eight paths into one of three evolutionary categories: direct, indirect, and combined. The direct group encompasses paths in which dN is directly related to tumour prevalence (paths 1 and 2). The indirect group includes paths in which dN is related to body size but not tumour prevalence (paths 3, 4, and 5). Finally, the combined group represents paths in which dN is related to both body size and tumour prevalence (paths 6, 7, and 8) (Figure 3A, Supplementary Table S4).

Across all dN genes, we observed a significant association between the evolutionary category and interaction of the dN effect direction and tumour type 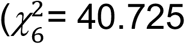, P < 0.001, N=1698) (Figure 3D). Indirect paths were significantly less common for dN-positive genes for both benign and malignant tumour prevalence (P < 0.001 and P < 0.001, respectively. In contrast, indirect paths (P = 0.001) and combined paths (P = 0.003) were significantly more common for dN-negative genes influencing benign tumour prevalence. Indirect and combined paths were also the most common among dN-negative genes that influence malignant tumour prevalence.

Finally, we found significant differences in the path prevalence between benign and malignant tumours for dN-negative 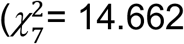, P = 0.041, N=1276, Figure 3B), dS-positive 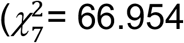, P < 0.001, N=2281, Figure 3C), and dS-negative 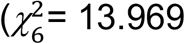, P = 0.030, N=77, Figure 3C) genes. For dS-positive genes, paths 3 and 8 were significantly associated with benign disease (P = 0.003 and P < 0.001 respectively, Figure 3C, 3E, 3G) while path 5 was significantly associated with malignant disease (P < 0.001, Figure 3C, 3F).

Paths 5 and 8, which both comprise causal relationships from dS to dN and tumour prevalence, dN to body size, and body size to tumour prevalence, were prominent among dS-positive genes. Both paths feature negative dN effects, implying that increased non-synonymous evolution is associated with reductions in body size, and in the case of path 8 a reduction in tumour prevalence. Similarly, both paths also include a positive effect of dS on dN, consistent with the nearly neutral expectation that a higher background rate of synonymous evolution increases the opportunity for non-synonymous change ^26^ (Figure 3F & G). However, path 5 captures a counteracting influence of dS: increases in dS are associated with a reduction in body size but an increase in tumour prevalence, highlighting a diametric effect of dS (Figure 3F). For example, we find that *WRN* is associated with path 5. In humans, null mutations in *WRN* cause Werner syndrome, which is characterised by premature aging and stunted growth, and also an exceptionally high incidence of sarcomas ^36^. While commonly associated with nonsense mutations ^36^, high-frequency synonymous substitutions have also been shown in patients with Werner syndrome ^37^.

Taken as a whole, these results demonstrate that genes that increase tumour prevalence tend to act directly or in combination with body size (or associated phenotypes). Conversely, genes that reduce tumour prevalence are more likely to act via body size, either indirectly or in combination with a direct effect. This reveals a broad evolutionary principle: reduced tumour prevalence is often achieved not through gene changes that target tumour suppression directly, but through genes that have shaped the evolution of body size itself.

### Species and genes that defy evolutionary expectations

To identify species with exceptionally high or low tumour prevalence, we compared the observed benign and malignant tumour prevalence of each species to the predicted values obtained from the full model (see Methods). Species with predictions that diverged substantially from observed data, exhibit exceptional tumour prevalence, or, in other words, are evolutionary outliers ^9^.

We found that nearly all species (105 out of 109) had a predicted benign or malignant tumour prevalence that differed by more than two standard-deviations from their observed prevalence in at least one gene (see Methods, Supplementary Table S5). Among the 99 species that had a predicted benign tumour prevalence that differed from the observed, we found that the dominant pattern was a lower-than-expected benign tumour prevalence, which occurred in 90 out of 99 species (Figure 4A). For instance, we found that *Columba livia* (Rock pigeon) and *Leucopsar rothschildi* (Bali myna) had a lower-than-expected benign tumour prevalence in > 40% of genes while *Ovis canadensis* (Bighorn sheep) had a lower-than-expected benign tumour prevalence in ~ 24% of genes. We found a similar pattern for malignant tumours in which 101 out of 105 species had a predicted malignant tumour prevalence that differed from their observed prevalence in at least one gene, and 88 out of 101 species had a lower-than-expected malignant prevalence (Figure 4B). For instance, *Momotus momota* (Amazonian motmot) had a lower-than-expected malignant tumour prevalence in ~ 46% of genes while *Aepyceros melampus* (Impala) had a lower-than-expected malignant tumour prevalence in > 30% of genes.

**Figure 4.**
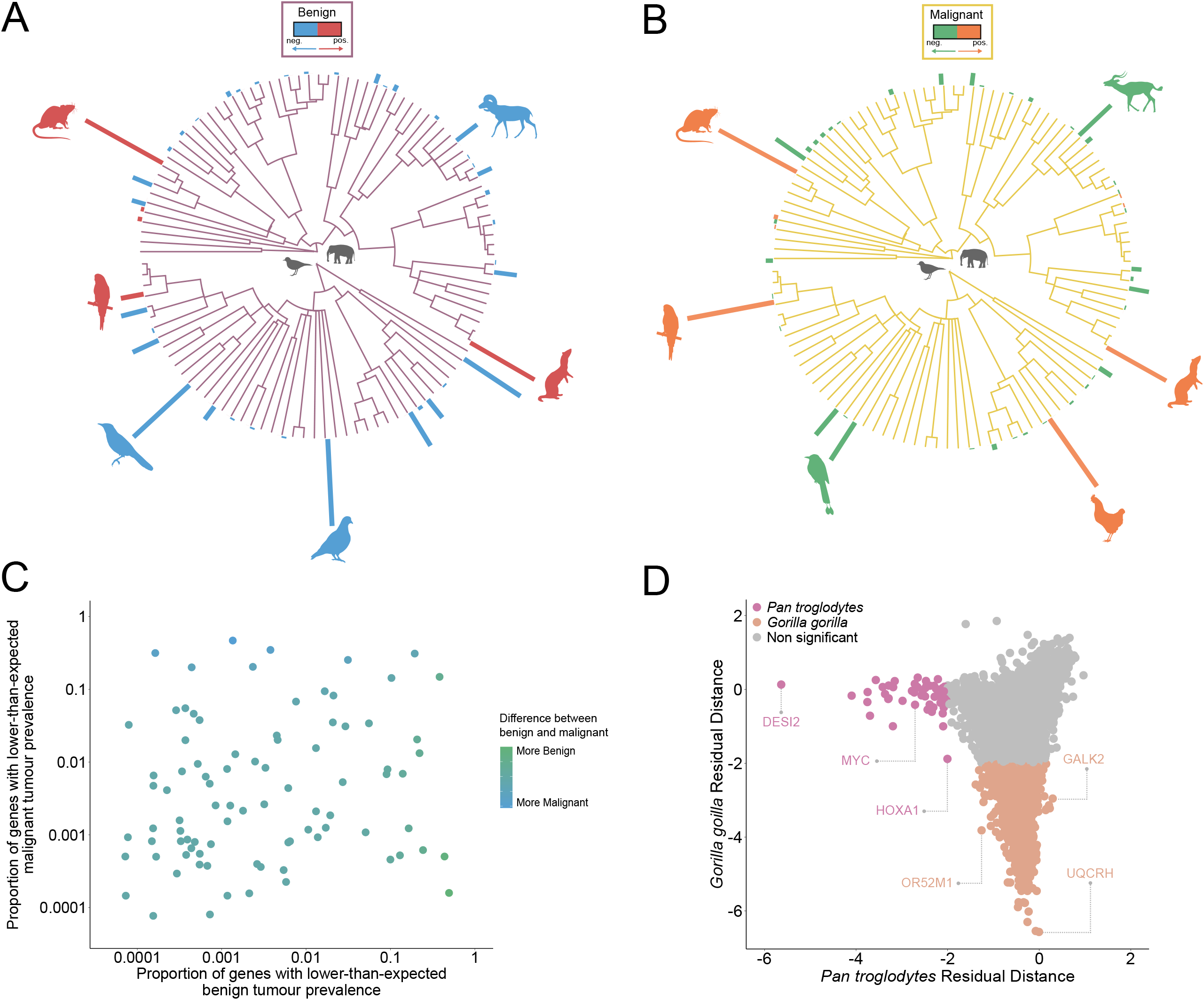
Evolutionary exceptions. **A**-**B**) The height of the bars at the tips of the phylogenetic trees corresponds to the proportion of genes in which the predicted **A**) benign or **B**) malignant tumour prevalence is significantly higher (red or orange) or lower (blue or green) than the observed. Species with a larger proportion of genes exhibiting lower- or higher-than-expected tumour prevalence represent stronger evolutionary outliers. **C**) Scatterplot indicating the proportion of genes in each species that exhibit lower-than-expected benign and malignant tumour prevalence, respectively. There is no association (P = 0.106) between the species-specific proportion of genes with lower-than-expected malignant and benign tumour prevalence. The colour of the points corresponds to the difference between the proportion of genes with lower-than-expected benign or malignant tumour prevalence. **D**) The malignant residual distance of each gene for *Pan troglodytes* (Chimpanzee) and *Gorilla gorilla (*Gorilla). Points in pink and terracotta correspond to genes in which *Pan troglodytes* and *Gorilla gorilla* have significantly less malignant tumour prevalence than would be expected. We find no overlap in significant genes between *Pan troglodytes* and *Gorilla gorilla* and no genes in which either species has a significantly higher prevalence of malignant disease than would be expected.

Despite the tendency for species to exhibit lower than expected tumour prevalence, benign and malignant deviations were not associated at the species level (P = 0.106, Figure 4C). Species with an exceptionally low benign tumour prevalence across many genes did not necessarily show a similarly low malignant tumour prevalence, indicating that benign and malignant tumours are shaped by distinct evolutionary pressures. For instance, *C. livia* had a lower-than-expected benign tumour prevalence in ~49% of genes but a lower-than-expected malignant tumour prevalence in < 1% genes. Similarly, *A. melampus*, had a lower-than-expected benign tumour prevalence in < 1% of genes but a lower-than-expected malignant tumour prevalence in > 30% of genes.

While the majority of the exceptional species had a lower-than-expected tumour prevalence, we found that a subset of species had a higher-than-expected tumour prevalence. Notably, *Rattus norvegicus* (brown rat) and *Mustela putorius* (ferret) had a higher-than-expected prevalence of benign tumours across 81% and 68% of genes, respectively (Figure 4A). Moreover, *Melopsittacus undulatus* (Budgerigar) and *Gallus gallus* (red junglefowl) had a higher-than-expected prevalence of malignant tumours across 91% and 85% of genes, respectively (Figure 4B). Interestingly, while *G. gallus* had a higher-than-expected prevalence of malignant tumours in over 80% of genes, it had a lower-than-expected prevalence of benign tumours in ~ 1% of genes, again, highlighting how benign and malignant growth can, and do, diverge within the same species.

Finally, we examined gene-level patterns of exceptional malignant tumour prevalence in our closest relatives, *Pan troglodytes* (chimpanzee) and *Gorilla gorilla* (gorilla) (Figure 4D). Consistent with the broader trends observed across birds and mammals (Figure 4B), all exceptional genes in both *P. troglodytes* and *G. gorilla* were associated with a lower-than-expected malignant tumour prevalence. Notably, *P. troglodytes* and *G. gorilla* did not share any of these exceptional genes (Figure 4D). This suggests that the genetic determinants associated with an exceptional reduction in tumour prevalence may be lineage-specific, even among closely related species. For instance, *P. troglodytes* exhibited lower-than-expected malignant tumour prevalence for *MYC* (a major known oncogene in human disease), highlighting how natural selection shapes key cancer-associated genes to drive lineage-specific patterns of tumour suppression (Extended Figure 3).

### Future directions

Taken as a whole, our results exemplify genetic pointillism at work – adaptive alterations, each small in isolation but powerful in aggregation, shaping tumour prevalence across millions of years of evolutionary change.

While we have focused on the effect of synonymous and non-synonymous alterations, our evolutionary framework can be used as a blueprint for future work seeking to link disease prevalence to rates of molecular evolution more broadly. Future studies considering the effect of additional molecular alterations, such as variation in the rate of copy number evolution ^38^ and methylation status ^39^, will be key to further understanding how natural selection shapes genome evolution more broadly and, in turn, its effects on tumour prevalence. Similarly, the same causal modelling framework used to disentangle the effect of body size, dN, or dS on tumour prevalence (Figure 3), presents an exciting opportunity to unravel how resistance mechanisms evolve across different molecular levels e.g. mutational, structural, and transcriptional variation. Relatedly, incorporating additional high-quality genomes and phenotypic datasets from future VGP releases will allow us to capture more subtle evolutionary signal and other clade-specific factors.

While comparative studies provide crucial insight into the evolutionary dynamics that shape tumour prevalence, species-specific mechanistic studies are still needed to understand why species such as *G. gallus* are particularly vulnerable to the scourge of cancer and yet its close relative *Pavo cristatus* (Indian peafowl) remains unscathed (Figure 4B). Similarly, future mechanistic investigations are needed to understand the key molecular differences between benign and malignant growths to identify cancer specific molecular vulnerabilities.

## Conclusion

The availability of high-quality genomic data ^17^, combined with phylogenetic statistical methods ^20,26^, provides an unprecedented opportunity to formally test genotype-phenotype relationships at a scale and resolution that has never been previously possible. Within individual species, we highlight the divergent evolutionary dynamics between benign and malignant tumours and identify two target groups of species to understand cancer susceptibility and mechanisms of cancer resistance. Throughout the genome, we demonstrate that body size plays a key, and previously underappreciated, role in mediating tumour suppression. Finally, at the individual gene level, we show that natural selection has been a precise and exacting sculptor counteracting the deleterious effect of nearly neutral processes.

## Supporting information

VGP Phase I Manuscript Extended Author List

## Extended Figure Legends

**Extended Figure 1:**
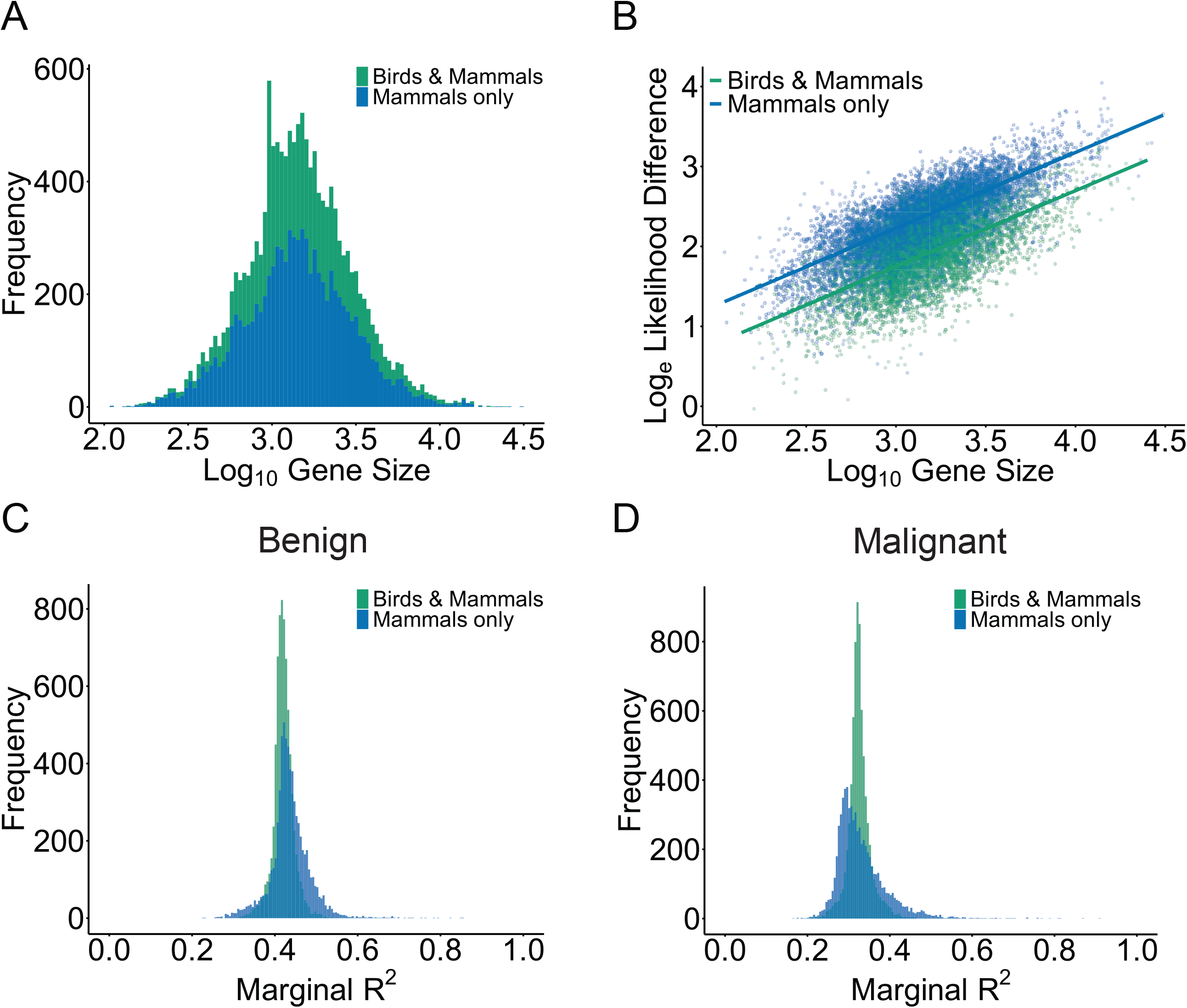
Linking to human disease. **A**-**D**) A volcano plot of the effect size of dN or dS on benign and malignant tumour prevalence. **E**-**F**) The proportion of overlapping significant **E**) dN-associated or **F**) dS-associated genes within each curated cancer dataset (COSMIC, IntOGen, OncoKB) stratified by whether the gene is associated with benign, malignant tumour prevalence, or both. **E**) Significantly more dN-associated genes are associated with malignant prevalence (COSMIC 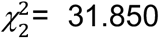 P < 0.001, N=120; IntOGen 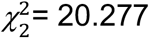, P < 0.001, N=101; OncoKB 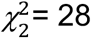, P < 0.001, N=206). Across all three datasets, malignant-associated genes comprise the largest overlap among significant dN-associated genes. **F**) Significantly more dS-associated genes are associated with benign prevalence (COSMIC 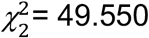 P < 0.001, N=160; IntOGen 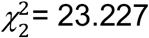, P < 0.001, N=132; OncoKB 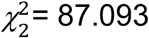, P < 0.001, N=247). Across all three datasets, benign-associated genes comprise the largest overlap among significant dS-associated genes.

**Extended Figure 2:**
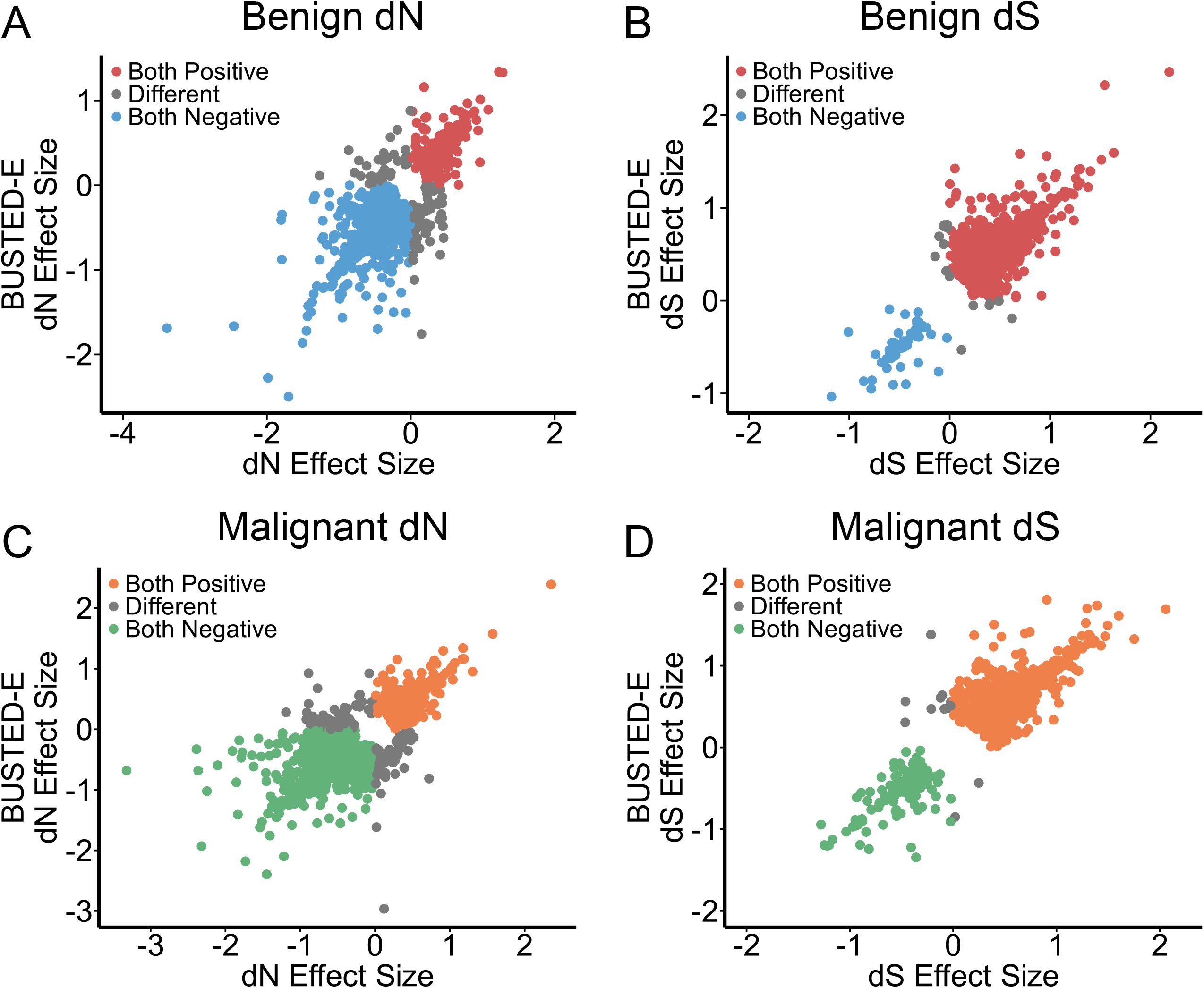
Conditional independencies and path analysis results. **A**) A schematic of the 11 different proposed causal relations (paths) that were tested between dS, dN, body size, and tumour prevalence. The direction of the arrow indicates the direction of the causal relationship. The background colour indicates whether the path is direct (orange; paths 1-2), indirect (green; paths 3-5), or combined (pink; paths 6-8). Paths P9 – P11 failed the conditional independence criterion and were not retained for causal modelling (see Methods). **B**) The proportion of 445 randomly selected genes with non-significant dN or dS stratified by the best fitting path. Among the eight retained paths, path 3 was the most common and accounts for ~ 50% of genes. **C)** The proportion of genes with a significant dN and dS association stratified by the best fitting path and coloured according to the dN direction and tumour type. Positive and negative dN associations are shown separately for benign and malignant tumour prevalence, highlighting differences in the distribution of causal paths among significant genes.

**Extended Figure 3:**
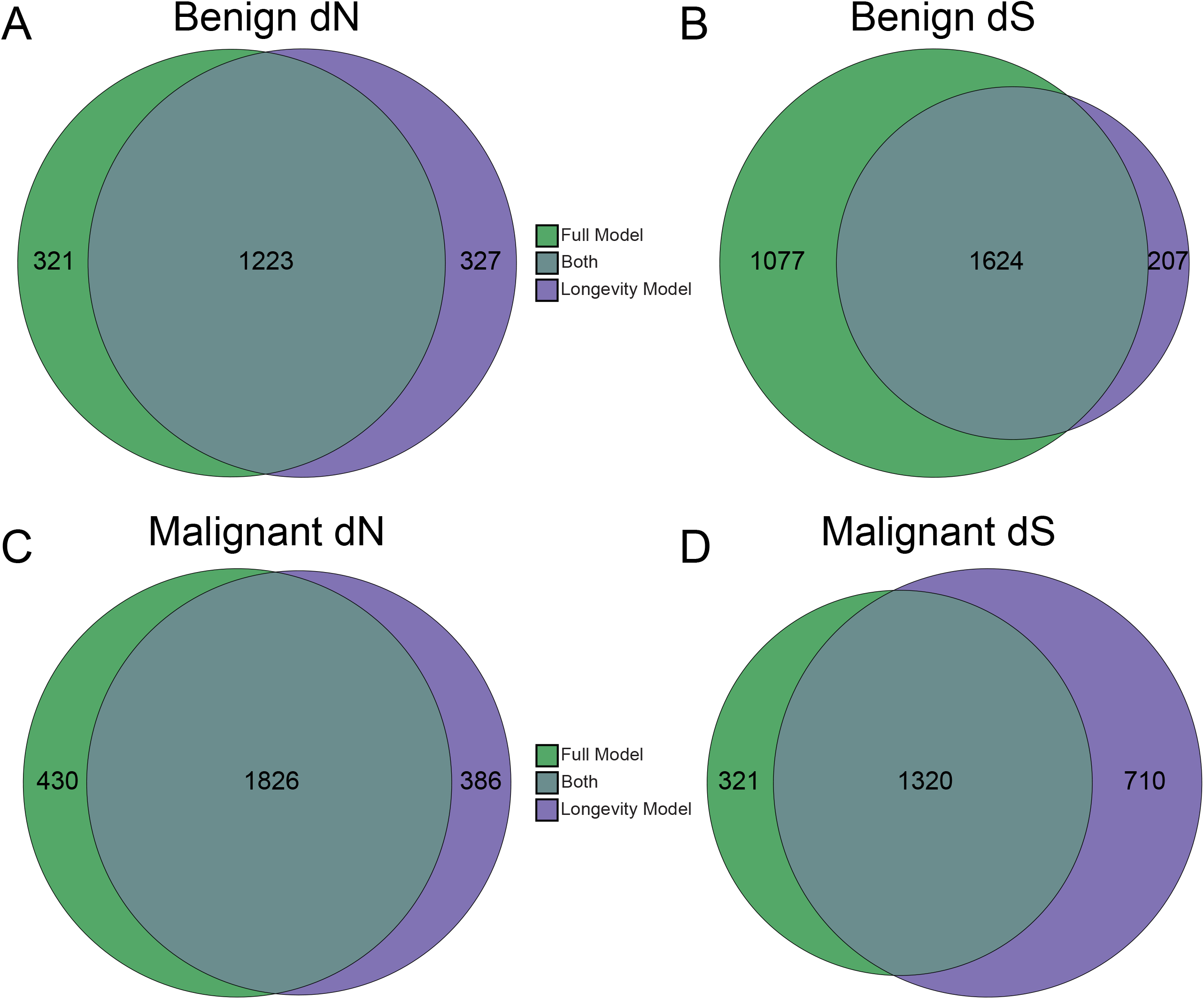
Evolutionary landscape of cross-species amino acid variation at functionally significant sites in *MYC*. Maximum-likelihood phylogeny of 130 vertebrate species (IQ-TREE 2, LG+G+F). Each column represents a high-impact *MYC* variant ranked by a weighted severity score (0.40 × normalized Grantham distance + 0.40 × SIFT severity + 0.20 × entropy), with one representative substitution shown per position. Cell colours denote amino acid states relative to human: alternate allele tracked across species (dark teal), other substitutions (salmon), identical to human (white), and gap (grey). Mutation labels are annotated by source database; COSMIC (lavender). Tree branches are coloured by major vertebrate groups (mammals (blue), birds (orange)). The right panel summarizes per-site metrics, including Grantham physicochemical distance (0–215), SIFT-derived severity (1 − SIFT score, 0–1), Shannon entropy (0–1), and alternate allele frequency across non-human species. Sites are ordered by decreasing weighted severity score. For visualization purposes, three outlier species with ~80% of sites differing from human were excluded.

## Supplementary Figure Legends

**Supplementary Figure 1:**
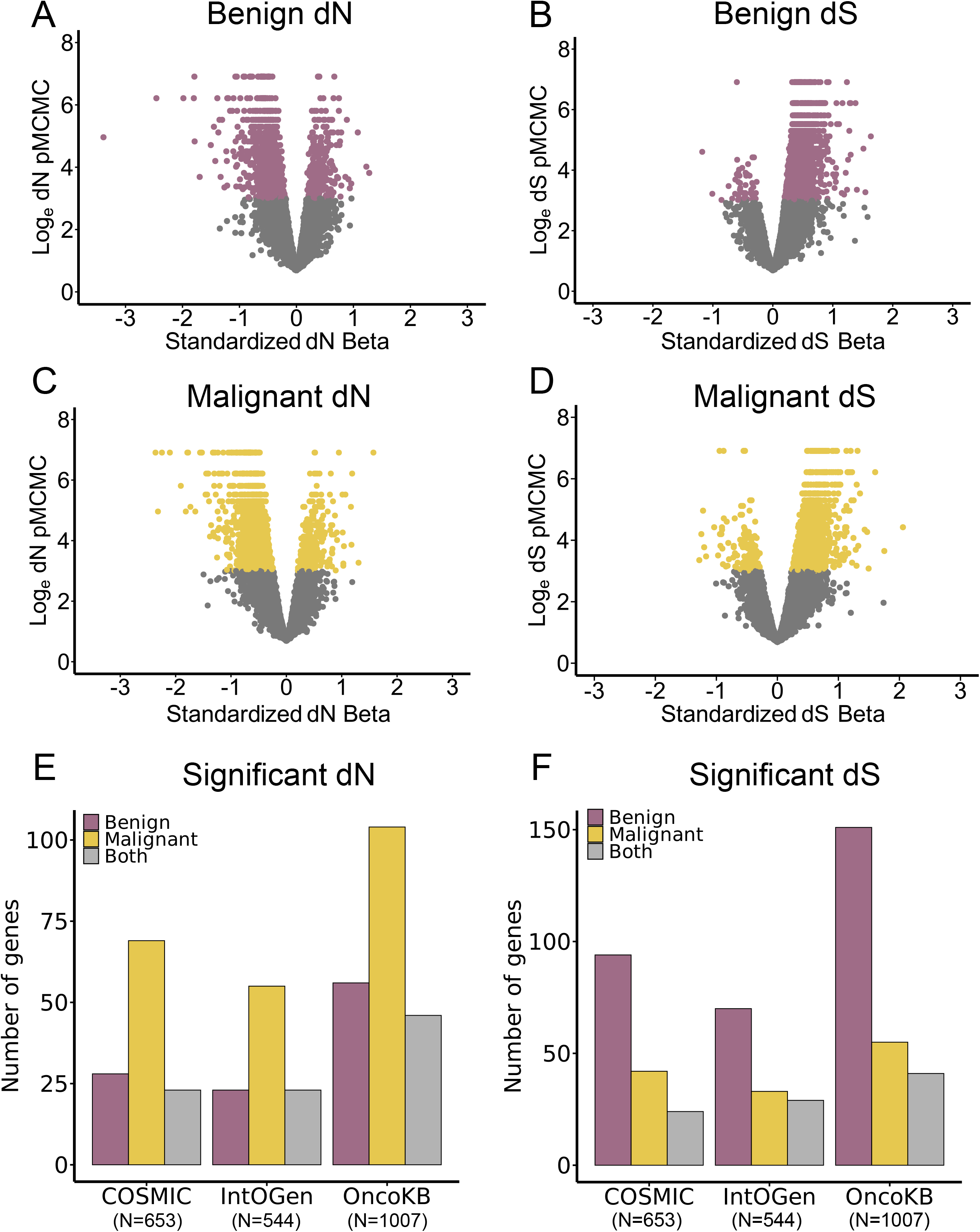
Assembly contiguity and gene conservation in Birds and Mammals. **A**) Distribution of scaffold N50 values for Birds (green) and Mammals (blue). **B**) Distribution of average gene identity scores for genes lifted from human using Miniprot. Identity scores represent the proportion of aligned amino acid residues that are identical between the lifted ortholog and the human reference sequence. **C**) Distribution of average positive substitution scores for genes lifted from human using Miniprot. Positive substitution scores represent the proportion of aligned residues that are identical or biochemically similar (conservative substitutions).

**Supplementary Figure 2:**
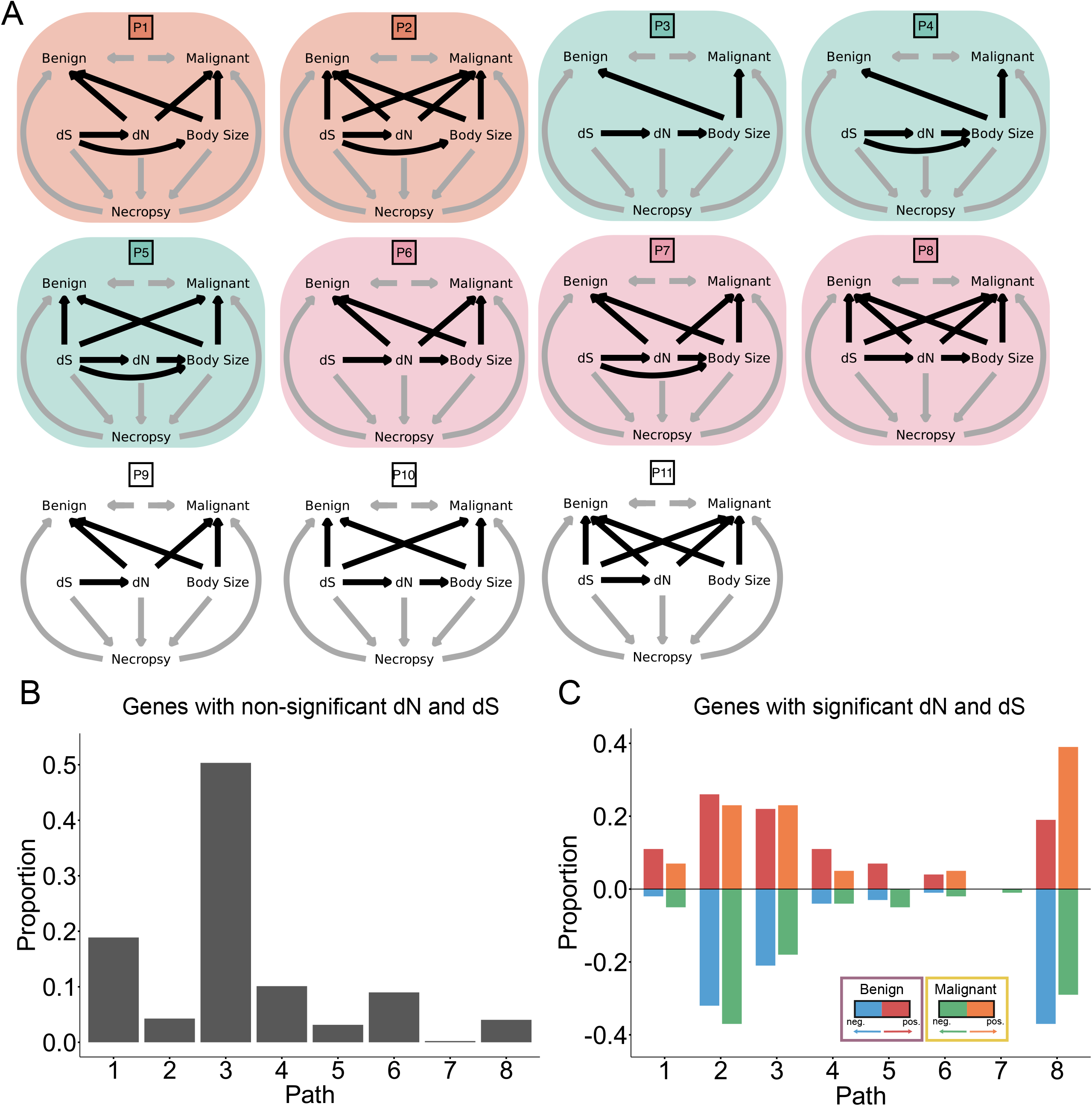
Model performance. In all cases, multiple sequence alignments (MSAs) containing birds and mammals are shown in green and mammals only are shown in blue. **A**) A histogram of gene size, measured as the number of nucleotides, stratified by the breadth of species in the alignment. **B**) A plot of the log-likelihood difference between fitting a single rate class per branch with no site-to-site variation compared to a full adaptive rate class model. The log-likelihood difference is positively associated with gene size in MSAs containing birds and mammals (p < 0.001) and mammals only (p < 0.001). **C**-**D**) Histograms of the marginal R^2^ values stratified by the number of classes for **C**) benign and **D**) malignant tumour prevalence (see Methods).

**Supplementary Figure 3:**
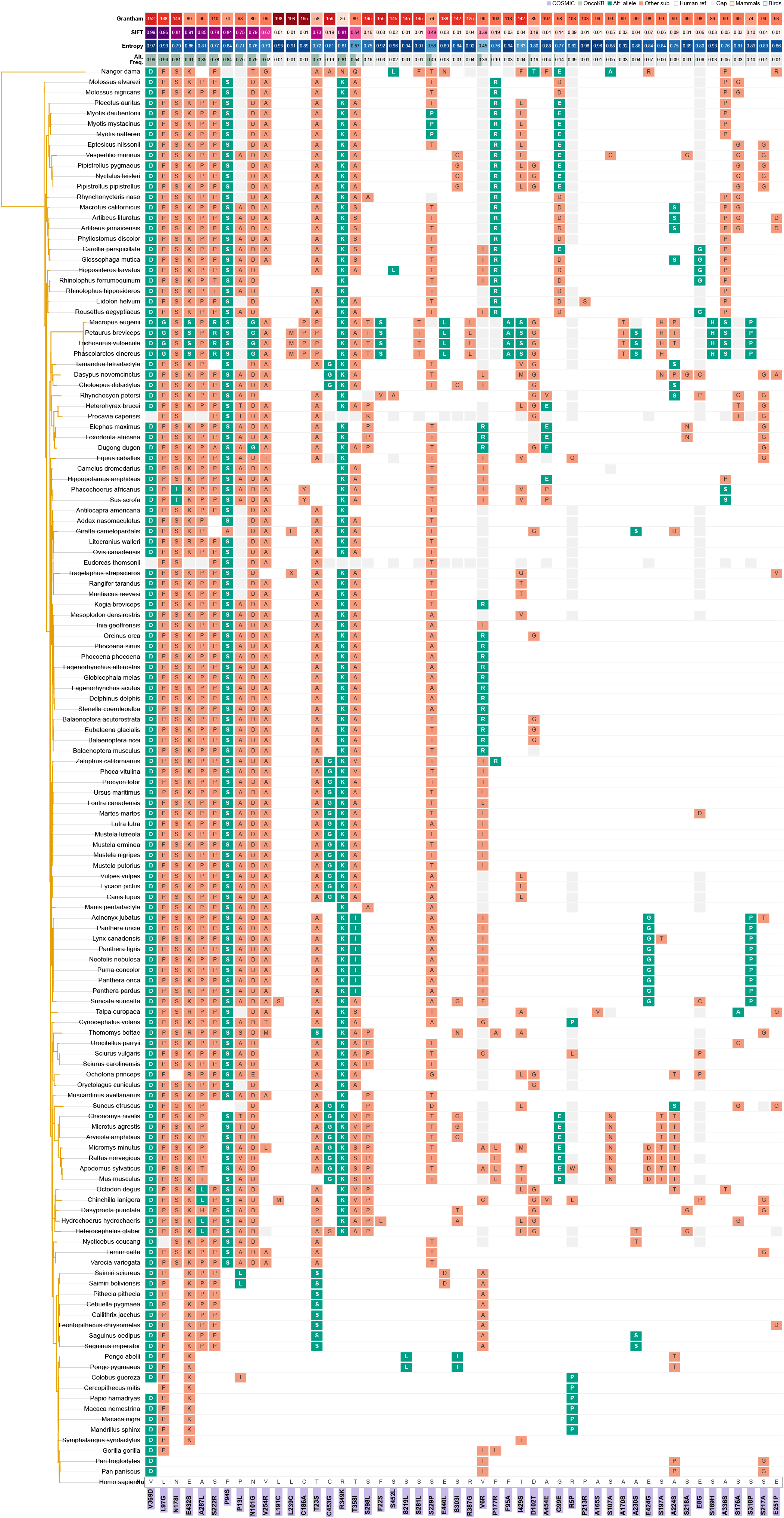
Evaluating the effect of alignment errors. Estimated dN and dS effect sizes on benign or malignant tumour prevalence with and without BUSTED-E filtering. **A-B**) Genes with a significant positive or negative association between dN or dS and benign tumour prevalence with and without BUSTED-E filtering are shown in red and blue respectively. **C-D**) Genes with a significant positive or negative association between dN or dS and malignant tumour prevalence with and without BUSTED-E filtering are shown in orange and green respectively. In all cases, genes with a significant dN or dS effect with and without BUSTED-E filtering but opposing directionality are shown in grey (see Methods). The strong concordance between effect size estimates before and after filtering indicates that the primary results are robust to potential alignment errors.

**Supplementary Figure 4:**
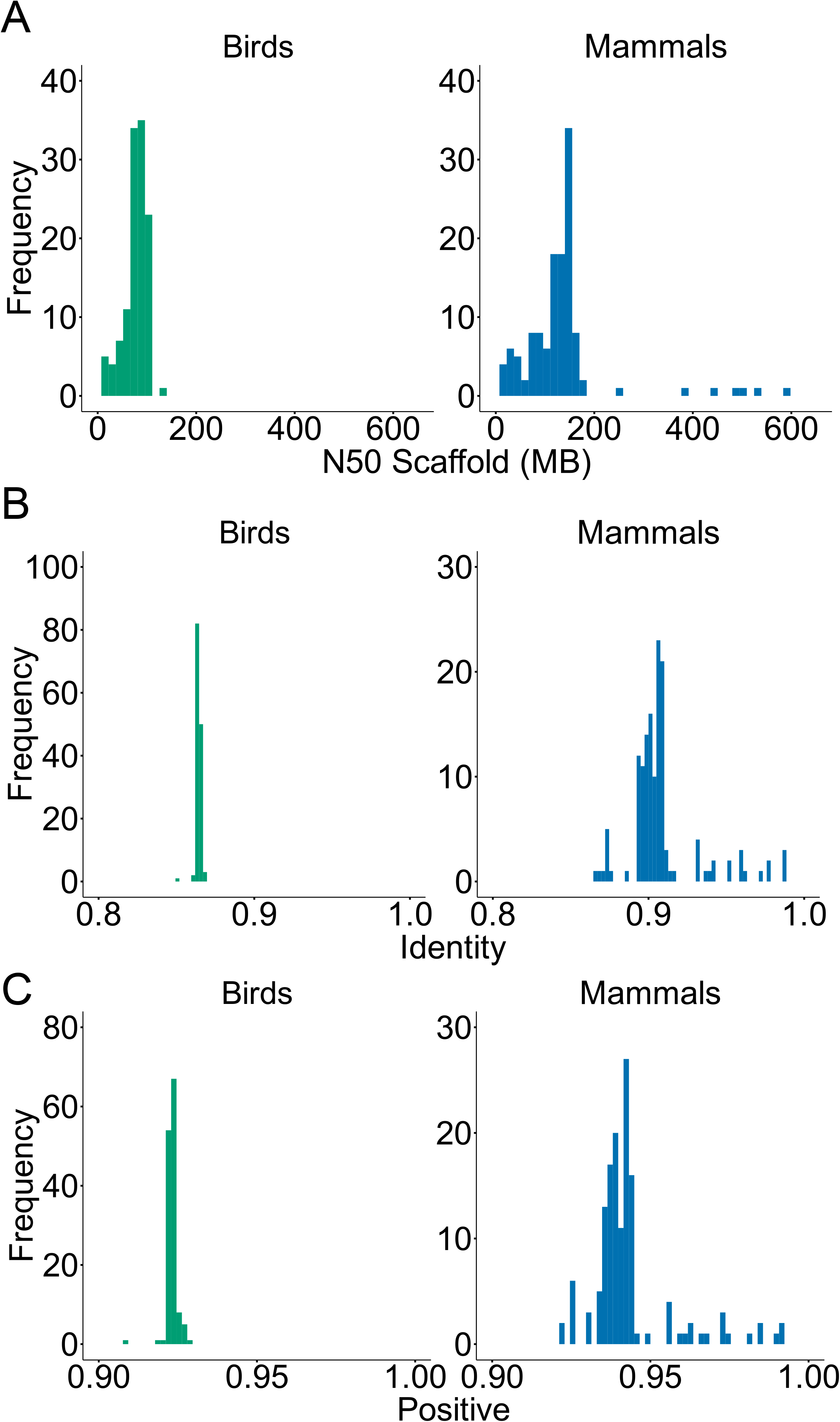
Controlling for species longevity. **A**-**D**) Comparison of significant genes identified using the full model and secondary model that included species longevity as an additional covariate (see Methods). Qualitatively similar patterns were observed between models with and without longevity included as an additional covariate, ‘longevity model’ and ‘full model’ respectively. Numbers indicate genes identified exclusively by the full model (green), exclusively by the longevity model (purple), or shared between both models (grey). The substantial overlap between models indicates that primary results are robust to the inclusion of species longevity.

## Methods

### Data and code availability

Benign and malignant tumour prevalence data for each species are available from Compton et al ^2^. As described by Compton et al, all necropsies were performed by specialist veterinary pathologists and neoplasms were identified by board-certified pathologists. Body mass data are available from Cooney and Thomas ^40^. Genomes are available from NCBI ^41^ (Supplementary Table S1). A phylogenetic tree from timetree.org was used to quantify molecular rate variation ^42^ (downloaded February 2024, see Quantifying molecular rate variation). A phylogenetic tree from Compton et al. ^2^ was used to model tumour prevalence (see Multivariate Phylogenetic Generalized Linear Mixed Models). Longevity data were collected from The Animal Ageing and Longevity Database (AnAge) ^43^. The final dataset of birds and mammals consisted of 109 species, 66 mammals and 43 birds (see below for details), spanning 14,189 individual genes (Supplementary Table S3). All necessary code is available at: https://github.com/george-butler/genome_wide_cancer_dynamics

### Building multiple sequence alignments (MSAs)

Among the initial 169 species of birds and mammals with paired tumour prevalence and body size data, 117 had corresponding genome assemblies available in the NCBI database ^41^ (Supplementary Table S1). When multiple assemblies were available for a given species, we selected the most recent genome, prioritizing long-read-based assemblies when available. A further 163 bird and mammal VGP genomes were also included to improve rate estimation. Similarly, a subset of amphibian genomes was also included to act as an outgroup for each alignment (see Multivariate Phylogenetic Generalized Linear Mixed Models). The final data for MSA construction consists of 280 species of birds and mammals.

From UniProt ^22^, we curated 20,421 reviewed human genes (Supplementary Table S2) as candidate orthologs for cross-species genomic analysis. Gene models were lifted over to the selected genomes using Miniprot ^21^, and custom scripts were applied to retain the best gene model per species based on alignment coverage and sequence identity. Gene models with ≥75% identity and coverage were kept, with more relaxed thresholds (≥40%) applied to the outgroup taxa. The resulting one-to-one orthologs were processed through a multiple sequence alignment and filtering pipeline adapted from OrthoMAM v12 ^23^ to ensure alignment quality and phylogenetic suitability. Genes represented in fewer than 50 species or lacking an amphibian outgroup were excluded from downstream analyses, leaving a final of 15,124 individual MSAs.

### Quantifying molecular rate variation

To detect variation in the rate of molecular evolution, an adaptive branch-site random effect model (aBSREL) was fitted in Hyphy for each gene ^24^. First, each alignment was filtered to remove species in which over 75% of the sites were missing. If over 75% of the sites were missing within the amphibian outgroup, the gene was removed from all further analyses. Next, for each gene, the aBSREL model was fitted using the time tree of life ^42^, trimmed to match the species within the alignment, and allowing for double and triple hits within a given site. A total of 256 species in the alignment matched species within the time tree of life (187 long read assemblies and 69 short read assemblies, with a subset of 112 species with matched cancer data). Once fitted, separate synonymous and non-synonymous rate-scaled phylogenies were generated, with branch lengths scaled to the branch specific rate of synonymous and non-synonymous evolution respectively. The two trees were rooted on the amphibian outgroups, which were then removed. Finally, branches were summed from root to tip for each species to obtain the total amount of synonymous and non-synonymous evolution, henceforth referred to as dS and dN respectively ^20^. This process was repeated independently for each gene.

### Multivariate Phylogenetic Generalized Linear Mixed Models (MPGLMMS)

Benign and malignant tumour prevalence was modelled throughout using multivariate phylogenetic generalized linear mixed models (MPGLMMs) fitted in a Bayesian Markov Chain Monte Carlo (MCMC) framework using the MCMCglmm R package ^19^. Specifically, benign and malignant tumour prevalence was estimated jointly for each gene with separate intercepts for birds and mammals. For genes where only a single class of species was present, e.g. only mammals, a single intercept was fitted. Phylogeny was included as a random effect in every model to account for the shared ancestry. The log_e_ total number of necropsies, log_e_ body mass, log_e_ dN, and log_e_ dS were included as standardized fixed effects, henceforth referred to as the full model (outlined below). That is, each independent variable was transformed to have a mean of 0 and a standard deviation of 1. Additionally, a second model, henceforth referred to as the longevity model, was also fitted with species longevity included as an additional covariate (outlined below). For genes where both birds and mammals were present, covariates were standardized on a per-class basis. At least five species needed to be present for a given class to be included in the model. Three species of marsupial (*Acrobates pygmaeus* (Feathertail glider), *Petaurus breviceps* (Pygmy sugar glider), *Phascolarctos cinereus* (Koala)) were removed prior to tumour modelling due to known differences in the immune system of marsupials compared to other mammals. A single slope was estimated for each dependent variable across all species. The final dataset consisted of 14,189 converged genes spanning 109 species of birds and mammals (Supplementary Table S3).

#### Full model

*Benign & Malignancy = Class + Log (Necropsy) + Log(Body mass) + Log(dN) + Log(dS) + Phylogeny*

#### Longevity model

*Benign & Malignancy = Class + Log (Necropsy) + Log(Body mass) + Log(dN) + Log(dS) + Log(Longevity) + Phylogeny*

#### MCMC conditions

All MCMC chains were run for 1×10^7^ iterations. The first 9×10^6^ iterations were discarded as burn-in, and the chain was sampled at every 1,000^th^ iteration. Models were considered to be converged if each of the fixed effects in the model had an effective sample size > 500. MPGLMMS were fitted with a Poisson link to account for the error structure in the benign and malignancy count data. MCMCglmm automatically accounts for overdispersion in count data. We used default priors 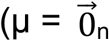 and V = **I**_n_ x 10^8^, where 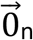 is the zero vector and **I**_n_ is the identity matrix in which n is equal to the number of fixed effects in the model) for the fixed effects and multivariate parameter-expanded priors (V = **I**_2_*2, ν = 2, 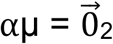, and ∝V = **I**_2_*25^2^, where 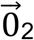 is the two dimensional zero vector and is the **I**_2_ two dimensional identity matrix) for the phylogenetic random effects ^44^.

#### Assessing significance and quantifying the variation explained by the model (R^2^)

Regression parameter significance was assessed by the proportion of the posterior distribution that crosses zero (P_x_), where P_x_ < 0.05 is considered to be significantly different from 0. The variation explained by the model was measured by calculating the pseudo-R^2^ for each gene. Specifically, univariate models were fitted separately for benign and malignant tumour prevalence and then the marginal and conditional R^2^ values were calculated using the method outlined by Nakagawa et al ^45^.

### Accounting for alignment errors

To control for potential alignment errors, we used the BUSTED-E model in Hyphy to preprocess each MSA ^46^. The BUSTED-E model was run with 5 starting points and with double and triple hits allowed within a given site. In turn, the filtered MSAs were then used to re-run the aBSREL model (see *Quantifying molecular rate variation*) and the subsequent MPGLMM analysis (see *Multivariate Phylogenetic Generalized Linear Mixed Models*).

We found a strong association between the estimated dN effect size with and without the BUSTED-E filtering for both benign (β = 0.796, P < 0.001, N=2132, R^2^ = 0.623, Supplementary Figure 3A) and malignant (β = 0.749, P < 0.001, N=2861, R^2^ = 0.577, Supplementary Figure 3C) tumour prevalence. Similarly, we found a strong association between the estimated dS effect size with and without BUSTED-E filtering for both benign (β = 0.855, P < 0.001, N=3332, R^2^ = 0.635, Supplementary Figure 3B) and malignant (β = 0.930, P < 0.001, N=2112, R^2^ = 0.799, Supplementary Figure 3D) tumour prevalence. The strong concordance in effect size with and without filtering reaffirm the quality of the genomes and subsequent MSAs and further support the robustness of our results.

### Linking to human disease

A total of 758 genes were obtained from the Cancer Gene Census (CGC) catalogue within the Catalogue Of Somatic Mutations in Cancer (COSMIC) database ^27^, of which 653 overlapped with the 14,189 genes in our dataset (www.cancer.sanger.ac.uk/census). From the OncoKB™ Cancer Gene List ^29^ 1192 genes were downloaded, including 1007 that overlapped with our dataset (www.oncokb.org/cancer-genes). Additionally, 633 genes were retrieved from the mutational cancer driver gene list in the Integrative Onco Genomics (IntOGen) database ^28^, of which 544 overlapped with our dataset (www.intogen.org). All three curated cancer gene datasets were downloaded on 6 June 2025.

### Causal inference modelling

Causal inference was conducted via a Phylogenetic Bayesian Structural Equation model (PhyBaSE) as outlined by von Hardenberg and Gonzalez-Voyer ^33^. Briefly, causal models were fit through a 3-stage process: defining causal models, testing conditional independencies, and causal inference via structural equation models

#### Defining causal models

A total of 11 different causal paths were proposed and defined as Directed Acyclic Graphs (DAG) (Extended Figure 2A). Across the 11 paths, causal relationships were set between the number of necropsies to benign and malignant tumour prevalence, from dS, dN, and body mass to the number of necropsies, from dS to dN, and from body mass to benign and malignant tumour prevalence. The covariance was also estimated between benign and malignant tumour prevalence. The remaining causal relationships between dS, dN, body mass, and benign and malignant tumour prevalence varied across all possible 11 paths.

#### Testing conditional independencies

Next, the implied conditional independencies in each of the 11 causal paths were tested as outlined in von Hardenberg and Gonzalez-Voyer ^47^. Specifically, each model was fitted as an MPGLMM in MCMCglmm ^19^. All MCMC chains were run for 1×10^7^ iterations. The first 9×10^6^ iterations were discarded as burn-in, and the chain was sampled at every 1,000^th^ iteration. Models were considered to be converged if each of the fixed effects in the model had an effective sample size > 500. We used default priors 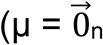 and V = **I**_n_ x 10^8^, where 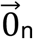 is the zero vector and **I**_n_ is the identity matrix in which n is equal to the number of fixed effects in the model) for the fixed effects and multivariate parameter-expanded priors (V = **I**_m_*m, ν = m, 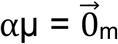, and ∝V = **I**_m_*25^2^, where 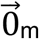 is the m dimensional zero vector and is the **I**_m_ m dimensional identity matrix in which m is equal to the number of dependent variables in the model) for the phylogenetic random effects.

A model was considered conditionally independent if 0 was included in the 95% credibility interval for all covariates in the model. The 11 different causal paths were fitted across the 5180 genes with a significant dN or dS covariate in the full model, and a random sample of 500 genes with a non-significant dN and dS covariate in the full model (total N = 5680). A path was considered to be suitable for causal inference if the path was conditionally independent in over 50% of the genes that were tested. Of the 11 paths that were tested, 8 were suitable for causal inference modelling.

#### Causal inference via structural equation models

Causal inference models were fitted across the 8 eligible paths for each gene. The causal inference models were fitted as an MPGLMM in MCMCglmm ^19^. All MCMC chains were run for 1×10^7^ iterations. The first 9×10^6^ iterations were discarded as burn-in, and the chain was sampled at every 1,000^th^ iteration. Models were considered to be converged if each of the fixed effects in the model had an effective sample size > 500. We used default priors 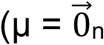 and V = **I**_n_ x 10^8^, where 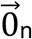 is the zero vector and **I**_n_ is the identity matrix in which n is equal to the number of fixed effects in the model) for the fixed effects and multivariate parameter-expanded priors (V = **I**_m_*m, ν = m, 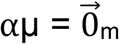, and ∝V = **I**_m_*25^2^, where 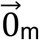 is the m dimensional zero vector and is the **I**_m_ m dimensional identity matrix in which m is equal to the number of dependent variables in the model) for the phylogenetic random effects. The best-fitting path for each gene was selected based on the lowest Deviance Information Criterion (DIC) (Supplementary Table S4).

### Identifying birds and mammals with exceptional tumour prevalence

We identified birds and mammals with exceptional tumour prevalence (either benign, malignant, or both) by comparing the species observed tumour prevalence to their expected level given their body size, dN, and dS. As outlined above (full model), MPGLMMs were fitted with benign and malignant tumour prevalence dependent on the number of records, body size, dN, and dS. The number of MCMC iterations, burn-in, sampling frequency, and prior specification were kept the same.

The fitted model was then used to predict benign or malignant tumour prevalence for each of the species within a given gene alignment. The predictions were made conditional on the estimates of the fixed and random effects. For each gene, a species was considered to have an exceptional tumour prevalence if the standardized residual distance was > 2 (Supplementary Table S5).

## Acknowledgements

We thank Alex Ostrovsky for assistance in accessing genomic data. This work was carried out at the Advanced Research Computing at Hopkins (ARCH) core facility (rockfish.jhu.edu), which is supported by the National Science Foundation (NSF) grant number OAC1920103. G.B is supported by the Prostate Cancer Foundation and the Patrick C. Walsh Prostate Cancer Research Fund. S.R.A is supported by the Prostate Cancer Foundation. M.C.S is supported, in part, by NIH awards U24CA284167, U24HG010263, and U41HG006620, NSF award 2419522, and the Lustgarten Foundation award 90101412. CV is supported by a Leverhulme Trust Leadership Award RL-2019-012. K.J.P is supported by the Prostate Cancer Foundation and National Cancer Institute, P50CA272391.

## Author Contributions

G.B and C.V designed the research. S.R, T.C, and M.C.S collected the genomes, developed the multiple sequence alignment (MSA) pipeline, and built the MSAs. G.B, and C.V developed the phylogenetic modelling, causal modelling, and evolutionary outlier analysis pipeline. G.B wrote the original draft of the manuscript. G.B, S.R, T.C, J.B, S.R.A, M.C.S, C.V, and K.J.P edited and revised the manuscript.

